# Individual natal assignment in highly migratory species: the genomic baseline and its application in loggerhead turtles

**DOI:** 10.64898/2026.05.06.723276

**Authors:** A. Luna-Ortiz, A. Barbanti, C. Pegueroles, F. A. Abreu-Grobois, P. Casale, D. Freggi, S. Giralt, E. Labastida-Estrada, R. Llera-Herrera, S. Machkour-M’Rabet, A. Marco, D. Margaritoulis, O. Turkozan, M. Pascual, C. Carreras

**Affiliations:** Departament de Genètica, Microbiologia i Estadística and IRBio, Universitat de Barcelona, Avinguda Diagonal, 643, 08028 Barcelona, Spain; Departament de Genètica i Microbiologia i Institut de Biomedicina i de Biotecnologia (IBB), Facultat de Biociències, Universitat Autònoma de Barcelona, Spain; Unidad Académica Mazatlán, Instituto de Ciencias del Mar y Limnología, Universidad Nacional Autónoma de México, Apartado Postal 811, Mazatlan, Sinaloa 82000 Mexico; Ethology Unit, Dept. of Biology, University of Pisa, Via A. Volta 6, 56126 Pisa, Italy; Lampedusa Turtle Rescue, c/o AMP, 92031 Lampedusa and Linosa Islands, AG, Italy; Fundación para la Conservación y la Recuperación de Animales Marinos (CRAM), 08820 El Prat de Llobregat, Spain; Laboratorio de Ecología Molecular y Conservación, Departamento de Conservación de la Biodiversidad, El Colegio de la Frontera Sur unidad Chetumal, 77014 Chetumal, Mexico; Estación Biológica de Doñana, CSIC, Av. Américo Vespucio s/n, 41092, Sevilla, Spain; ARCHELON, the Sea Turtle Protection Society of Greece, Solonos 113, Athens, Greece; Aydın Adnan Menderes University, Faculty of Science, Department of Biology, 09010 Aydın, Türkiye

**Keywords:** 2b-RAD, *Caretta caretta*, foraging grounds, migratory species, sea turtles, natal origin, population structure

## Abstract

1. Effective conservation of highly migratory species requires understanding genetic structure across breeding populations and access high□resolution markers capable of assigning individuals from mixed aggregates (e.g. bycatch or new nesting sites) to their natal origins. Genomic approaches provide unprecedented resolution but add methodological challenges; thus, it is essential to first build a genomic baseline from known breeding areas and then evaluate strategies for assigning unknown individuals.
2. To address this, we used 2b-RAD sequencing, a genomic reduction technique useful for degraded DNA, and loggerhead turtles as a case study. This species shows philopatric breeding, while juveniles and adults form mixed aggregations in foraging grounds.
3. Our results highlight the importance of building baselines that include all potential source populations contributing to mixed aggregations. We detected hierarchical genetic differentiation and high resolution and successfully assigned the natal origin of 124 unknown individuals from four Mediterranean foraging grounds. These grounds showed distinct source contributions, and comparisons with previous studies suggest possible temporal shifts in stock composition.
4. We provide a comprehensive genomic baseline for individual assignment of Altanto-Mediterranean loggerhead turtles of unknown natal origin and a general framework for identifying population-specific threats in highly migratory species.

## 1 INTRODUCTION

The wide spatial distribution, long lifespan, and use of multiple, dispersed habitats for development, foraging, and reproduction make highly migratory species particularly challenging to monitor and conserve. Movements among these habitats expose individuals to diverse anthropogenic threats across life stages, including fisheries interactions, pollution, habitat degradation, and vessel collisions (Cooke et al., 2024). Individuals from multiple breeding populations can share foraging areas without mating (Carreras et al., 2011). Consequently, a key requirement for effective management is the ability to reliably assign individuals at developmental and foraging sites to their natal origins. Conservation genetics should therefore extend beyond breeding populations to quantify the composition of mixed assemblages and help delineate migratory links between breeding and foraging habitats (Vela-Garcia et al., 2025).

Dispersal has traditionally been inferred from gene flow, uncovering population structure (Hohenlohe et al., 2021). However, directly identifying the breeding origin of individuals sampled away from natal areas is equally critical. This requires two complementary datasets: a baseline dataset comprising individuals of known origin, and a mixed (or unknown) dataset. First, a robust genomic baseline must be established using individuals from potential source populations. Second, unknown individuals must be genotyped and compared against this baseline to infer their natal origin. Integrating both aspects provides a comprehensive understanding of population structure and dispersal, which is essential for effective conservation planning (Labastida-Estrada & Machkour M’Rabet, 2024).

Sea turtles are emblematic highly migratory and endangered species, undertaking long-distance migrations between nesting and foraging areas that can span thousands of kilometres and more than a decade (Hays et al., 2002). Despite this, they exhibit natal philopatry, returning to their birthplace to reproduce, which generates detectable genetic structure (Clusa et al., 2018; Barbanti et al., 2025). Combining philopatry-driven genetic structure with migratory patterns has led to defining Regional Management Units (RMUs) as multiple genetically distinct populations sharing migration routes, foraging areas, and threats (Wallace et al., 2023). Understanding connectivity within and among RMUs is therefore central to conservation, particularly as individuals from different RMUs may co-occur in mixed foraging aggregations (Wallace et al., 2023).

Mixed stock analysis (MSA) estimates the proportion of individuals originating from different populations in a potential mixed aggregate (Grant et al., 1980). In sea turtles, MSA has primarily relied on mitochondrial DNA, revealing that Mediterranean foraging grounds host individuals from multiple RMUs (Mediterranean, North-West Atlantic and North-East Atlantic) (Vela-Garcia et al., 2025). However, the widespread sharing of mtDNA haplotypes among source populations results in large confidence intervals, limiting the precision of stock composition assessments (Carreras et al., 2006; Barbanti et al., 2019). Nuclear microsatellites enabled individual assignment (IA) in marine loggerhead turtles (Carreras et al., 2011) but constrained individual assignments to the RMU level, and left many individuals unassigned (Clusa et al., 2016).

Recent genomic advances now allow the use of thousands of markers, greatly enhancing resolution. In this context, 2b-RAD sequencing is particularly suitable for species with large genomes and degraded DNA (Barbanti et al., 2020). A recent 2b-RAD study has revealed unprecedented fine-scale population structure in Mediterranean nesting populations and identified distinct sub-regional groupings (SubRMUs) (Barbanti et al., 2025), demonstrating the potential of genomics to establish robust baselines for IA. However, the absence of Atlantic samples limits its application for assigning individuals found in Mediterranean foraging grounds. Moreover, genotyping and filtering procedures can substantially influence downstream analyses (Hemstrom et al., 2024), highlighting the need for standardized and reproducible approaches.

Our study aims to establish best practices for applying genomic baselines to assign individuals of unknown natal origin in highly migratory species. Using *Caretta caretta* as a model, we evaluate how genomic differentiation across RMUs and SubRMUs can improve individual assignment. We specifically how alternative genotyping strategies for both baseline and unknown datasets affect assignment outcomes. By integrating genomic data across breeding and foraging habitats, this work provides a framework to quantify population contributions to mixed aggregations and offers general guidelines for genomic individual assignment in conservation and management.

## 2 MATERIALS AND METHODS

### 2.1 Sampling

We built a baseline dataset with 274 individuals from 14 nesting areas (Table S1), representing three Regional Management Units (RMUs) known to contribute to Mediterranean foraging grounds (Figure 1). The dataset included 15 individuals from the North-West Atlantic RMU (NWA; 12 from Quintana Roo (QRO), Mexico, and 3 from Grand Cayman (CAY), Cayman Islands), 25 from the North-East Atlantic RMU (NEA; all from Boa Vista (BOA), Cabo Verde), and 234 previously published unrelated individuals from the Mediterranean (MED) RMU (Barbanti et al. 2025). Nesting-area samples were obtained from hatchlings or nesting females.

**Figure 1.**
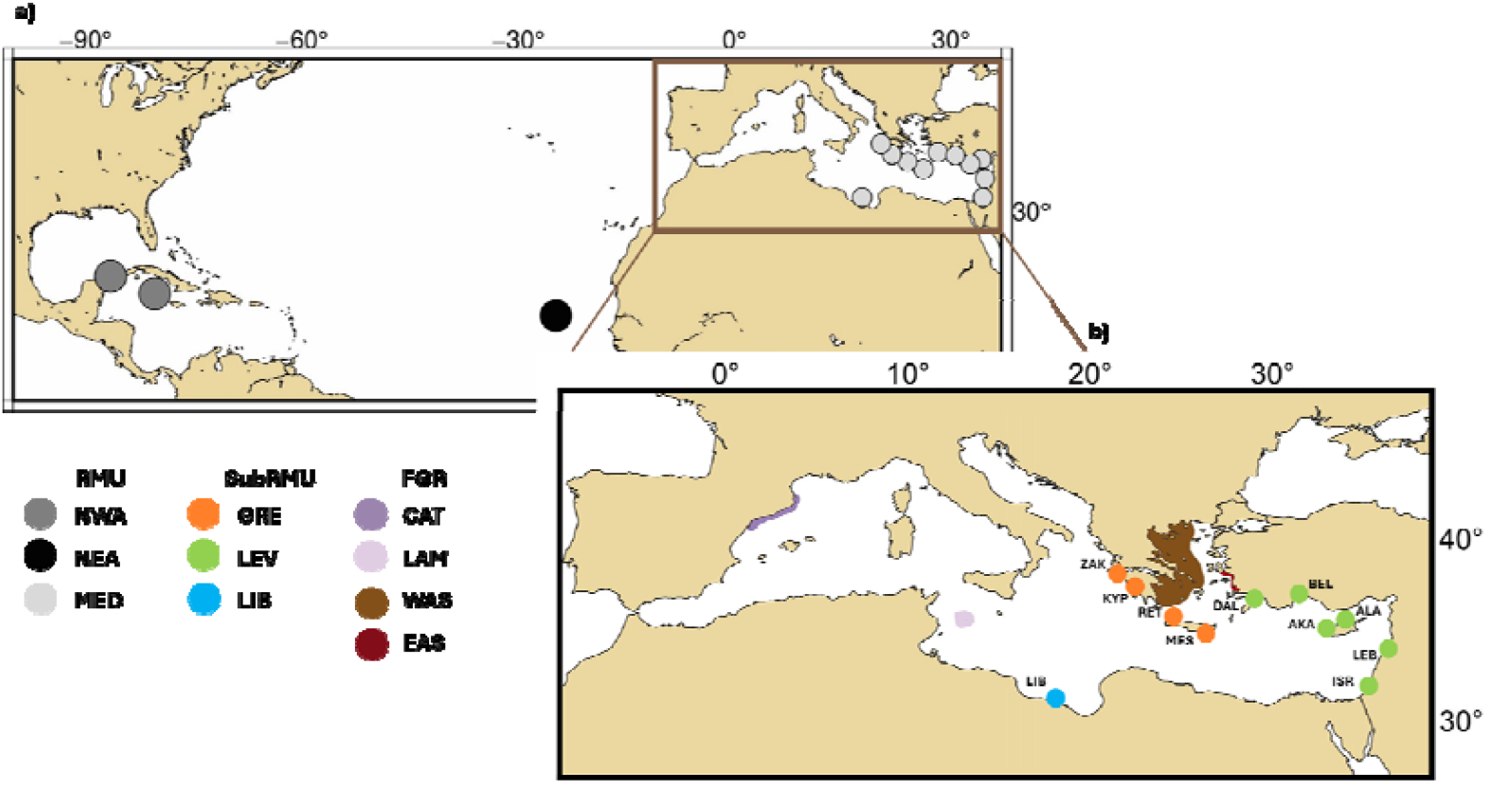
Map of nesting locations (baseline) and foraging grounds (unknown natal origin). **a)** Baseline samples were from North-West Atlantic (NWA), North-East Atlantic (NEA), and Mediterranean (MED) RMUs. **b)** Mediterranean locations were grouped in three SubRMUs: Libya (LIB); Levantine (LEV; Israel (ISR), Lebanon (LEB), Alagadi (ALA), Akamas (AKA), Belek (BEL), and Dalyan (DAL)); and Greece (GRE; Messara (MES), Rethymno (RET), Kyparissia (KYP), and Zakynthos (ZAK)). Coloured polygons indicate the sampled foraging areas of individuals of unknown origin: Catalan coast (CAT), Lampedusa (LAM), Western Aegean coast (WAS), and Eastern Aegean coast (EAS). Maps created with QGIS (https://qgis.org/).

Additionally, we analysed 124 individuals of unknown natal origin collected from four Mediterranean foraging grounds (FGRs) (Figure 1, Table S1): 32 from the Catalan coast (CAT), 38 from the central Mediterranean (Lampedusa area, LAM), 23 from the West Aegean coast (WAS), and 31 from the East Aegean coast (EAS). These samples were obtained from animals admitted to rescue centres. Skin samples were collected from dead or live animals, while blood samples were collected from living animals following established rescue-centre protocols. All samples were preserved in 96% ethanol and stored at -20°C until DNA extraction.

### Laboratory and genotyping protocols

Genomic DNA was extracted using the Puregen Kit (Qiagen), following the manufacturer’s instructions. Individual libraries were prepared for the 164 individuals sampled in this study (Table S2) as for the 234 previously published Mediterranean baseline individuals (Barbanti et al., 2025). Briefly, 180 ng of DNA were digested with AlfI, and ligated to base-selective adapters (5’-WN-3’) to reduce the number of loci while preserving genetic differentiation (Galià-Camps et al., 2022). Fragments were amplified, purified using magnetic beads and quantified using the Quant-iT™ PicoGreen™ dsDNA Assay Kit (Thermo Fisher Scientific). Libraries were pooled by index type, ensuring ∼180 ng of DNA per library, except for QRO samples for which DNA was purified from 1.8% agarose gels. Sequencing was performed at the Centre for Genomic Regulation using an Illumina HiSeq 2500 for single-index libraries and a NextSeq 500 for double-index libraries.

Raw 2b-RAD reads from new and published data (398 individuals) were processed using customized scripts (https://github.com/EvolutionaryGenetics-UB-CEAB). Sequences were trimmed to remove adaptors, cut to the same length (34 bp), and mapped to the loggerhead reference genome (GenBank accession GCA_023653815.1, Chang et al., 2023) using Hisat2-2.2.183 (Kim et al., 2019). Two genotyping strategies were implemented to evaluate the effect of dataset composition (Figure S1). First, all samples were genotyped simultaneously (*Joint Genotyping*) with BCFtools (Li, 2011), and resulting genotypes saved in a VCF file from which baseline and unknown samples were separated using the *--keep* function in VCFtools (Danecek et al., 2011). Second, baseline and unknown samples were genotyped independently (*Non-Joint Genotyping*) and independent VCF files generated for each group.

In both strategies, the baseline VCF file was filtered using VCFtools by removing genotypes supported by fewer than five reads, loci exceeding 50X mean depth (defined as 1.5 times the interquartile range), and SNPs with missing data in more than 30% of individuals (*--max-missing* 0.7). SNPs with minor allele frequency >0.002 were kept, removing singletons. The SNPs retained in the baseline were extracted from the unknown dataset using the *--snps* function in VCFtools. The strategy retaining the highest number of informative SNPs in the unknown dataset was selected for downstream analyses.

Additionally, to evaluate the effect of sample size under the *Non-Joint Genotyping* approach, we performed subsampling analyses of the unknown dataset. We generated subsets of 10, 50 and 75 individuals, each with 10 random replicates sampled without replacement. Each replicate was genotyped independently, and the number of SNPs shared with the baseline was quantified. Differences among subset sizes were assessed using a Kruskal-Wallis test followed by post-hoc Wilcoxon signed-rank tests in R.

### Baseline genomic differentiation

With the best genotyping strategy, we built the baseline including the 274 samples separated by sampling locality (named *Caretta Baseline*) and characterised its population structure. Genetic differentiation among RMUs was assessed through pairwise F_ST_ values (Weir & Cockerham 1984) grouping sampling localities by RMU (Table S1). Statistical significance was assessed with 999 permutations using the R package ‘StAMPP’ (Pembleton et al., 2013). We estimated pairwise F_ST_ values and significance among the three Mediterranean SubRMUs using the *MED Baseline* dataset restricted to Mediterranean individuals and the same SNPs as the *Caretta Baseline*, with localities grouped by SubRMU (Table S1). Finally, we estimated pairwise genetic differentiation between all localities with the *Caretta Baseline* dataset.

Additionally, we evaluated genomic differentiation using Discriminant Analysis of Principal Components (DAPC), implemented in ‘adegenet’ (Jombart, 2008), following Jombart and Collins (http://adegenet.r-forge.r-project.org/files/tutorial-dapc.pdf). The optimal number of principal components was selected with ‘x.val’, and the number of genetic clusters (K) using ‘find.cluster’. We extracted the density distribution of the first two discriminant axes for both baseline datasets (*Caretta* and *MED*).

#### Individual assignment of the samples of unknown natal origin

We used the R package ‘assignPOP’ version 1.1.4 (Chen et al., 2018) for individual assignment, following Marín-Capuz et al. (2025). Five predictive models (Support Vector Machine (SVM), Latent Dirichlet Allocation (LDA), NaiveBayes (NAIVE), decision Tree (TREE), and Random Forest (FOREST)) were evaluated. Monte Carlo cross-validation using ‘assign.MC’ was performed to cluster individuals into reference and test data sets. For each model, 30 iterations were run using training proportions of 0.5, 0.7, 0.9, and loci proportions of 0.1, 0.25, 0.5 and 1.

The 124 samples of the unknown dataset were assigned using the ‘assign.X’ function and the best-performing model following three strategies: 1) assignment against the *Caretta Baseline* grouped by RMU (NWA, NEA, and MED); 2) assignment against the *MED Baseline* grouped by SubRMU (LIB, LEV, and GRE); and 3) assignment against the *Caretta Baseline* considering five groups (NWA, NEA, LIB, LEV, and GRE), referred to as ‘all-at-once’ approach.

## RESULTS

### Strategies for building a genomic baseline

Overall, 4,678,552 mean reads were obtained per individual (SD±1,727,584), with a mean mapping success of 93.08% (SD±7.35), regardless of the library preparation strategy (Table S2). After filtering the baseline datasets, 6,586 SNPs were retained in both genotyping strategies (Figure S1). However, the two approaches differed in the number of SNPs recovered in the unknown dataset. The *Joint Genotyping* strategy recovered all polymorphic markers identified in the baseline, whereas the *Non-Joint Genotyping* strategy recovered only 88.2%. Subsampling analyses under the *Non-Joint Genotyping* approach (Figure S2) revealed a strong effect of sample size on SNP recovery, with a significant reduction in in the number of shared loci at small sample sizes (Kruskal-Wallis p<0.05, Wilcoxon signed rank test, p<0.05 for all cases). Consequently, we selected the *Joint Genotyping* strategy for all subsequent analyses.

### Genomic differentiation of the baseline

All pairwise F_ST_ values between RMUs were statistically significant (Table S3a) and DAPC analysis revealed three genetic clusters corresponding to the three RMUs (Figure S3a). The first discriminant function clearly separated individuals from the three RMUs, with the greatest differentiation observed for Mediterranean individuals (Figure S3b), whereas the second discriminant function distinguished between individuals from the two Atlantic RMUs (Figure S3c). SubRMUs pairwise F_ST_ values with the *MED Baseline* were all significant (Table S3b) and the DAPC analysis revealed three distinct genetic clusters LIB, LEV and GRE (Figure S3d). The first discriminant axis differentiated the three SubRMUs with some overlap, whereas the second discriminant axis separated LIB from the rest (Figures S3f and S3e). Consequently, both baselines showed good differentiation and discrimination potential. Finally, we evaluated the genetic differentiation at the locality level using the *Caretta Baseline* (Figure S4) and found significant genetic differentiation between all pairwise localities except for AKA-ALA and DAL-BEL (Table S3c).

### Individual assignments in foraging areas

Natal origins of individuals from foraging grounds (FGR) were inferred with ‘assignPOP’ using the SVM model having the highest accuracy (Figure S5). Each assignment was repeated 10 times, and mean assignment probabilities per individual were calculated. First, we assigned the FGR individuals at the RMU level using the *Caretta Baseline* (Figure 2). All FGR individuals were assigned to a RMU with a probability >0.75 (Figure 2b; Table S4). We chose this threshold after plotting the assignment probabilities of the FGR individuals with the three baseline strategies (RMU, SubRMU, and all-at-once) to detect break points in their distribution (Figure S6). Most FGR individuals (96.8%) were assigned to MED (Table S4), three were assigned to NWA (one from CAT and two from LAM), and one to NEA (from LAM). Consequently, LAM had the highest proportion of individuals from Atlantic origin (7.9%).

**Figure 2.**
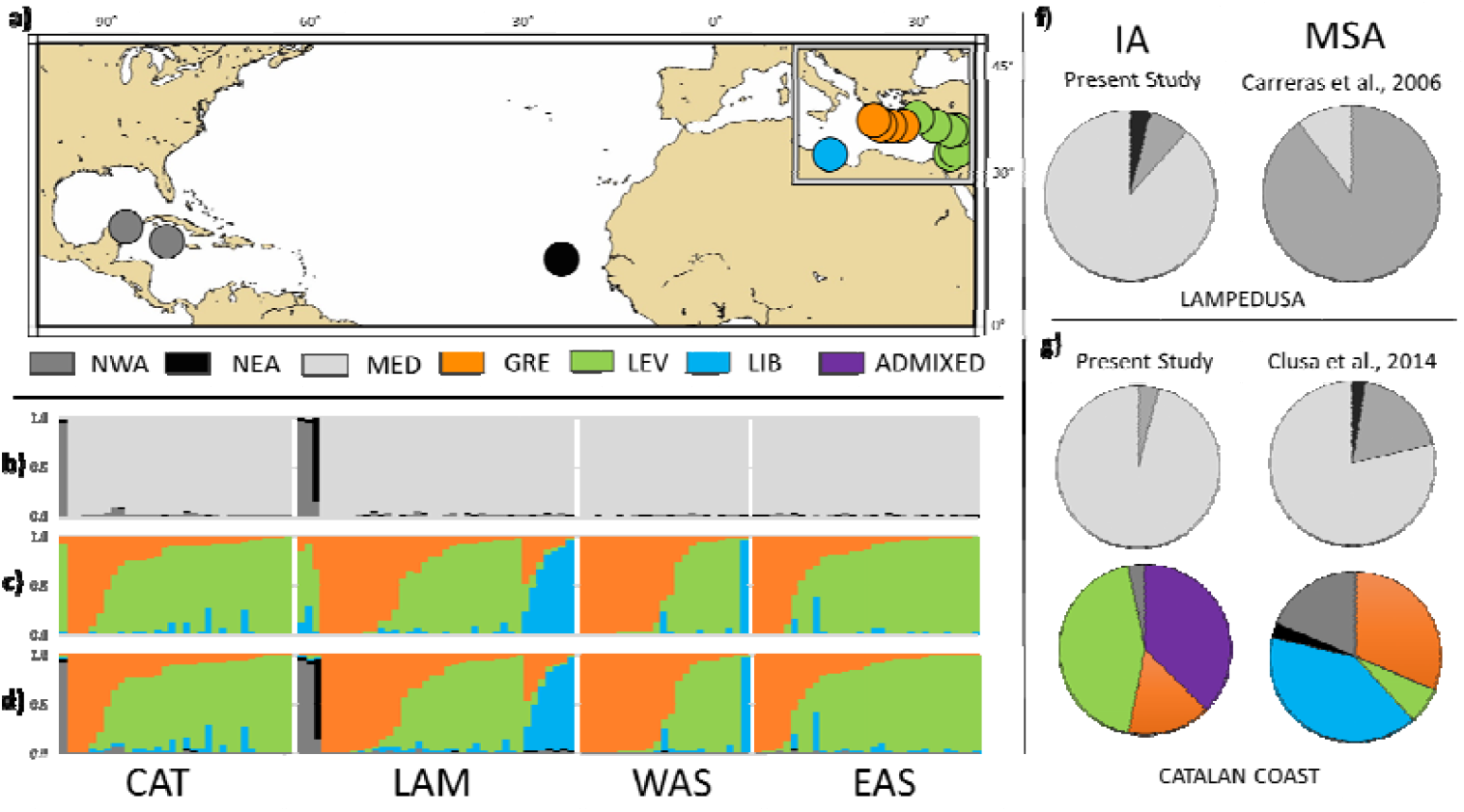
Results of the assignment strategies for individuals sampled in CAT, LAM, WAS and EAS foraging grounds (FGRs). **a)** nesting locations of the genomic baseline RMU coloured (North-West Atlantic: NWA, North-East Atlantic: NEA, and Mediterranean: MED), and SubRMUs coloured (Libya: LIB, Levantine: LEV and Greece: GRE). **b)** Individual assignment (IA) probabilities using the *Caretta Baseline* grouped at RMU. **c)** IA probabilities using the *MED Baseline* grouped at SubRMU. **d)** All-at-once IA probabilities using the *Caretta Baseline* grouped into five genetic groups (NWA, NEA, LIB, LEV, and GRE). The 124 FGRs individuals have the same order across panels. **f)** Proportion of individuals from Lampedusa (LAM) assigned to each RMU in this study and using Mixed Stock Analysis (MSA) in Carreras et al. (2006). **g)** Proportion of individuals from the Catalan Coast (CAT) assigned to RMU and with all-at-once approach in this study, and compared with MSA results from Clusa et al. (2014). Individuals with probabilities <0.75 to one group are considered admixed (purple). Maps created with QGIS (https://qgis.org/).

Second, FGR individuals were assigned using the *MED Baseline*. Assignments to a single SubRMU were considered successful with probabilities >0.75 (Table S5, Figure S6b), individuals with maximum assignment probability <0.75 are considered admixed. In total, 97 individuals were assigned to a single SubRMU (five to LIB, 61 to LEV, and 31 to GRE) and 27 individuals exhibited admixed origin (Figure 2c, Table S5). Of the four individuals previously assigned to an Atlantic origin, one was assigned to LEV, and the other three identified as admixed. This highlights that excluding potential source populations from the baseline leads to wrong assignments.

Third, FGR individuals were assigned using the all-at-once strategy (Figure 2d, Figure S6c). Three individuals were assigned to NWA and one to NEA, consistent with the RMU assignment, five were assigned to LIB, 57 to LEV, and 31 to GRE, while the remaining 27 were considered admixed. Aside from discrepancies caused by excluding Atlantic populations from the baseline, all other individuals presented similar assignment probabilities to the same SubRMU or considered admixed, except for individuals CC2006, CC2015 and CC2019 (Tables S5 and S6). Although individuals assigned to the Levantine SubRMU were most prevalent (46%), origin distributions varied by foraging ground (Figure 2d). Lampedusa presented the highest proportion assigned to Libya (10.5%), while WAS had the greatest proportion assigned to Greece (47.8%). The Catalan Coast (37.5%) and Lampedusa (21.1%) exhibited the highest frequency of admixed individuals (Table S6).

## DISCUSSION

Non-model and highly migratory species often exhibit complex life cycles and population structure, which can be challenging for population studies and for scientifically based management and conservation. In this study, we show that genomic techniques can avoid these shortcomings using marine turtles as a case study. First, we describe different hierarchical levels of genetic structure among nesting populations of loggerhead turtles across the Atlantic and Mediterranean. Second, we use this genomic information as a baseline to perform individual assignments for turtles sampled in Mediterranean foraging grounds relevant for its management and conservation. Finally, we provide guidelines for using genomic individual assignment in conservation and management of endangered migratory species.

The present study is the first demonstrating that RMUs (Wallace et al., 2023) are genomically fully supported, confirming a real intermediate level of genetic structuring in the species. Moreover, our reanalysis of Mediterranean nesting areas confirmed an additional SubRMU level (Barbanti et al., 2025). Overall, we show a hierarchical genetic organization of loggerhead turtles in at least three levels: populations, SubRMUs and RMUs. Unveiling different levels of population structure are fundamental in species management, and genomics is key in unveiling them. Gaps in sampling areas using the same methodology, such as Florida in NWA, should be considered in future studies as they may provide unused genetic groups (Silver-Gorges et al., 2026).

Our study shows that genotyping all the samples of unknown natal origin simultaneously with the baseline samples using the reference genome is needed (Figure 3). To optimise computational resources and reduce the size of the output file, genotyping softwares only provide information of polymorphic positions in relation to the reference genome (Li 2011). The number of recovered loci depends on sample size and genetic diversity. As shown in our simulations, genotyping small sets of unknown samples independently can substantially reduce markers and compromise assignment accuracy.

**Figure 3.**
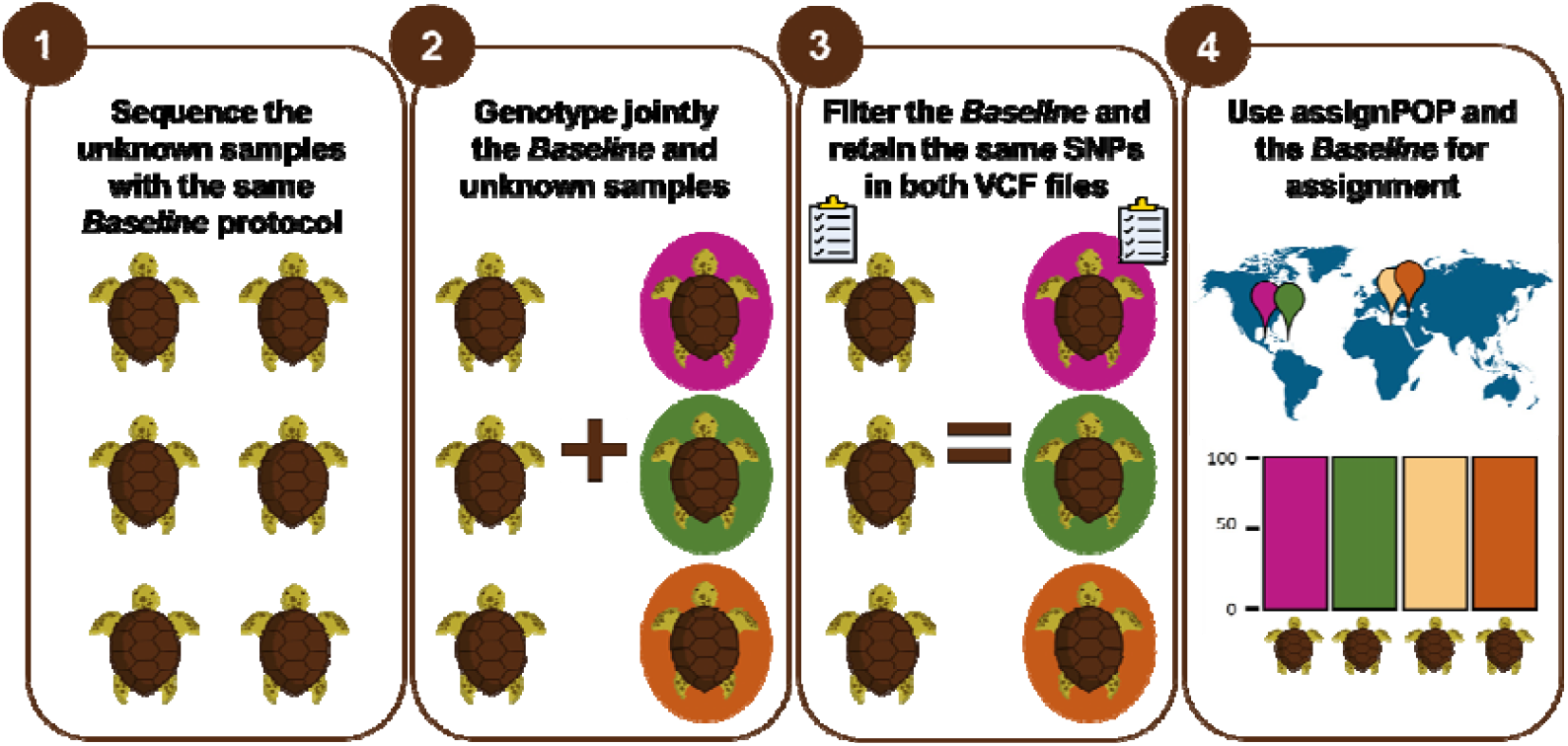
Pipeline for assigning individuals of unknown natal origin using a baseline.

A robust baseline is also essential for accurate assignment. Across all approaches, correct assignments occurred only when the baseline captured the genetic variation of the individual’s origin; otherwise, assignPop misassigned individuals with high confidence. This highlights the need for comprehensive baselines covering many nesting sites and hierarchical levels. Our genomic individual assignment outperformed mixed stock analysis, achieving full assignment thanks to more markers and a broader baseline, whereas MSA showed high rates of unassigned individuals (Clusa et al., 2016). Two valid strategies emerge for species with hierarchical genetic structure from our work: (1) a stepwise approach from RMU to SubRMU, removing individuals already assigned to other RMUs before finer analyses; and (2) an all-at-once approach that assigns individuals across both levels simultaneously. Both methods detected admixed individuals among SubRMUs, consistent with previous findings (Barbanti et al., 2025), suggesting these intermediate assignments reflect true admixture rather than methodological artifacts.

The composition of origins varied markedly among Mediterranean foraging areas, consistent with earlier mitochondrial MSA studies linking this heterogeneity to prevailing currents, although we detected a substantially lower proportion of Atlantic individuals (Clusa et al., 2014, Carreras et al., 2006). While comparisons should be made cautiously due to methodological differences and large MSA confidence intervals, these discrepancies likely reflect temporal shifts in foraging ground composition. A decline in Atlantic individuals may result from the recovery of Mediterranean nesting populations (Mazaris et al., 2017; Casale et al., 2018; Margaritoulis et al., 2025), increasing local recruitment and diluting Atlantic contributions. Changes in the Atlantic Meridional Overturning Circulation may also limit Atlantic turtles entering the Mediterranean (Ditlevsen & Ditlevsen, 2023).

In agreement with the hypothesis that migration from nesting to foraging grounds is shaped by Mediterranean current systems (Millot & Taupier-Letage 2004; Casale & Mariani 2014), the foraging areas of the Aegean Sea were composed of individuals from Greece and the Levantine region. Notably, the Aegean Sea showed subregional structuring: eastern areas were dominated by Levantine individuals while western areas had a higher contribution of Greek-origin individuals. Lampedusa exhibited a mixed composition from all three SubRMUs, with a slight Levantine proportion. Strikingly, the Catalan coast was dominated by Levantine individuals, with few Atlantic turtles and no Libyan contribution—contrasting sharply with earlier MSA estimates (Clusa et al., 2014). This discrepancy highlights the risk of interpreting MSA mean values without considering their associated error. The recent expansion of Turkish nesting populations (Casale et al., 2018), may explain the widespread dominance of Levantine juveniles, although increases in some Greek rookeries are also reported (Margaritoulis et al., 2025). Future studies using the same methodology across additional foraging areas will be essential to provide a better picture of the migration routes of juveniles in the Mediterranean.

### Conclusion

In this work, we propose a methodology to assign the natal origin of unknown individuals and guide genomic baseline design for highly migratory species (Fig 3). First, use the same library building methodology employed in the baseline in the unknown samples. Minor adjustments (e.g., indexing or pooling) do not affect genotyping results, demonstrating robustness across laboratories. Second, genotype new samples jointly with the baseline. Third, split the VCF into baseline and unknown files, filter the baseline, and generate a list of SNPs to retain in both datasets. Fourth, use assignPOP to determine the origin of the unknown individuals. For loggerhead turtles, we provide baseline sequences and a SNP panel for future assignments of unknown samples across the Atlanto-Mediterranean region. Moreover, our methodology is transferable to other non-model species and guide the generation of key information for managing species with complex life cycles.

## Acknowledgements

This work was supported by the projects MarGech (PID2020-118550RB) funded by MICIU/AEI/10.13039/501100011033, GenoMarTur (CNS2022-135205) funded by MICIU/AEI/10.13039/501100011033 and the European Union NextGenerationEU/PRTR), BlueDNA (PID2023-146307OB) funded by MICIU/AEI/10.13039/501100011033 and ERDF/EU and the contract: “Utilización de herramientas de secuenciación masiva como nuevos marcadores genéticos en tortuga boba (*Caretta caretta*), con el fin de probar su eficacia en la asignación de zonas de origen frente a los marcadores tradicionales, FB 04/2023 “, in the framework of the project “LIFE IP-PAF INTEMARES [LIFE15 IPE ES 012] “Gestión integrada, innovadora y participativa de la Red Natura 2000 en el medio marino español”, coordinated by the Biodiversity Foundation.. ALO, AB, CP, MP and CC are members of the research group SGR2021-01271 funded by the Generalitat de Catalunya. DM thanks the ARCHELON staff and volunteers for collecting the samples at the nesting beaches of Greece and at the ARCHELON Sea Turtle Rescue Centre in Glyfada. All the samples had the authorization of the corresponding country authorities. ALO dedicates this paper to Juan Luis Flores Colín.

## Data Availability

2b-RAD raw data can be found at the European Nucleotide Archive (ENA), for the Mediterranean nesting rookeries (project PRJEB77620, from Barbanti et al., 2025), for the Atlantic nesting rookeries (project PRJEB105674), and for the individuals sampled in four Mediterranean foraging grounds (project PRJEB105699). The name of the SNP loci used in the study and obtained by mapping the raw reads to the loggerhead genome (GenBank accession GCA_023653815.1, Chang et al., 2023) are available for future analyses as supplementary data (Baseline_SNPs.txt) as well as the genotypes of all individuals (Baseline.283indv.FGR.124indv.6598sites.vcf).

## Author Contributions

ALO, MP and CC designed the research. AAG, PC, DF, SG, ELE, SMM, AM, DM and OT provided samples. ALO, AB and RLH did the laboratory. ALO did the genomic data analyses with inputs from CP, MP and CC. ALO, MP and CC provided funding acquisition. ALO, MP and CC wrote the manuscript with inputs from all authors.

