## Supplementary Figures for "Individual natal assignment in highly migratory species: the genomic baseline and its application in loggerhead turtles"

**SUPPLEMENTARY MATERIAL**

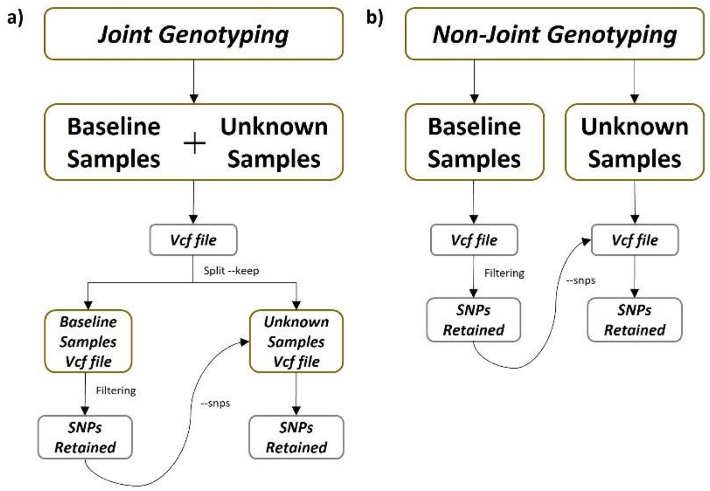


**Figure S1. Flowchart of the two genotyping strategies generating a list of common SNPs for comparison between individuals of the genomic baseline and unknown origin.** a) ***Joint Genotyping Strategy***, all samples are genotyped together. Once the VCF file is obtained, the baseline samples and unknown samples are separated into two VCF files. Filters are applied only to the baseline VCF file. As a result, a list of retained SNPs from the baseline is generated and used to extract the same SNPs from the VCF file of samples of unknown origin. b) ***Non-Joint Genotyping Strategy***, the mapped samples from the two groups are genotyped separately, one consisting of the baseline samples and the other of the samples of unknown origin. Once the VCF file of the baseline samples is obtained, the filters are applied, and a list of retained SNPs is created to be extracted from the VCF file of samples of unknown origin.

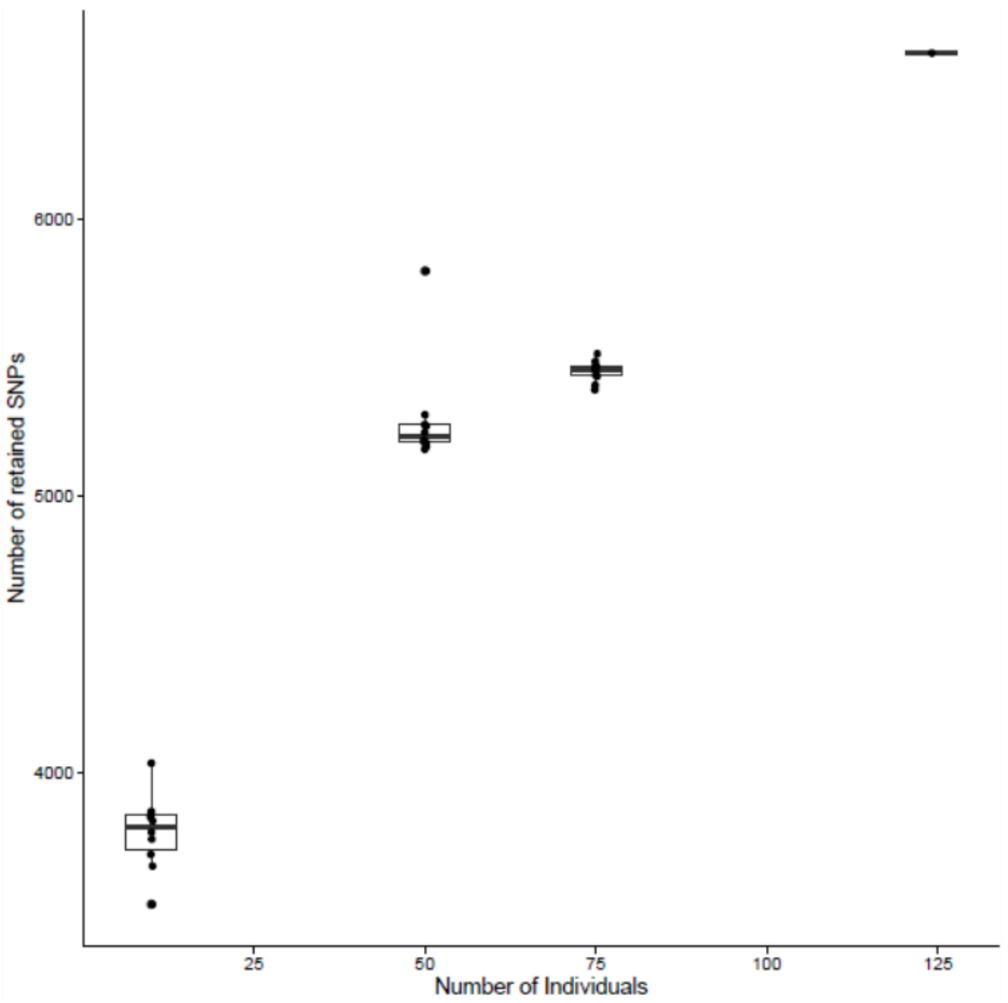


**Figure S2. Effect of unknown individuals sample size on SNP recovery.** Boxplot with the number of SNPs recovered on each one of the three different sample sizes tested (10, 50 and 75 individuals), and SNPs recovered on the whole dataset (124 individuals) in relation to the baseline. Each dot represents a different randomized list of individuals, drawn without replacement from the 124 individuals with unknown natal origin sampled in the Mediterranean foraging grounds. Ten randomized lists were generated for each of the three sample sizes.

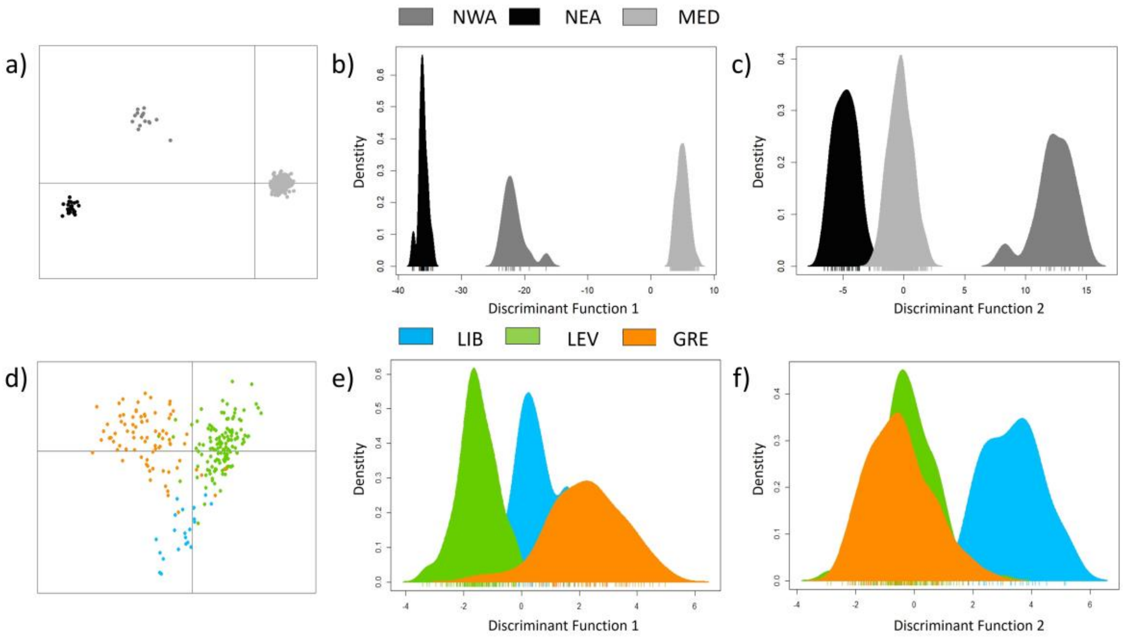


**Figure S3. Genetic structure of the baseline nesting locations for the loggerhead sea turtle (*Caretta caretta*) at the Regional Management Unit (RMU) and at the Sub Regional Management Unit (SubRMU) level.** After filtering, a panel of 6,5**86** SNPs was used to assess genomic differentiation at two levels. a) Discriminant analysis of Principal Components (DAPC) at RMU level with samples from the 14 nesting sites coloured according to their corresponding RMU (Table S1). b-c) RMU density plots along the two first discriminant functions. d) DAPC at SubRMU level with samples from the 11 Mediterranean nesting sites colour coded according to the three previously defined SubRMUs (Barbanti et al. 2025). e-f) SubRMU density plots along the two first discriminant function. Acronyms for RMU are NWA (North-West Atlantic), NEA (North-East Atlantic) and MED (Mediterranean). Acronyms for SubRMU are LIB (Libya), LEV (Levantine Region) and GRE (Greek Region). Note that each dot in the DAPC plots (a and d panels) represents an individual.

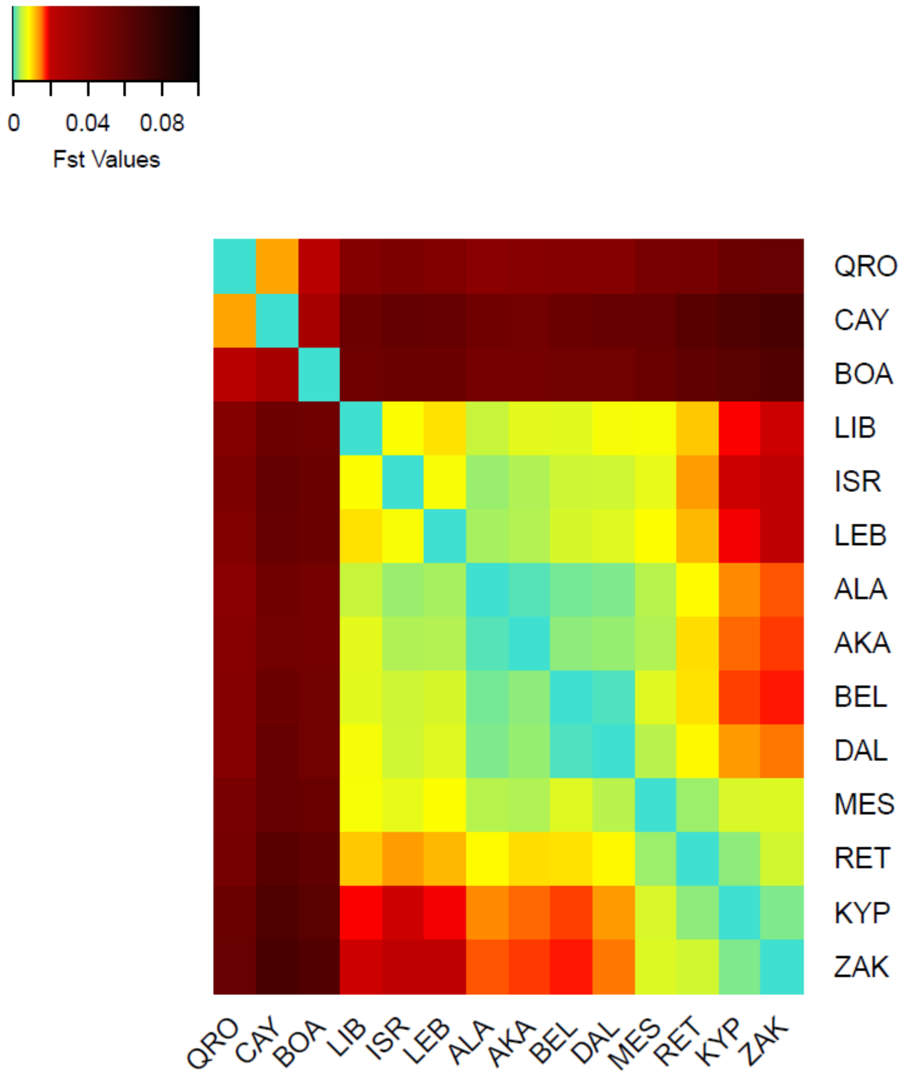


**Figure S4. Heatmap of the pairwise genetic differentiation ($F_{ST}$) between rookeries.** Note that the individuals of the *Caretta Baseline* dataset were separated by sampled nesting locations (Table S1).

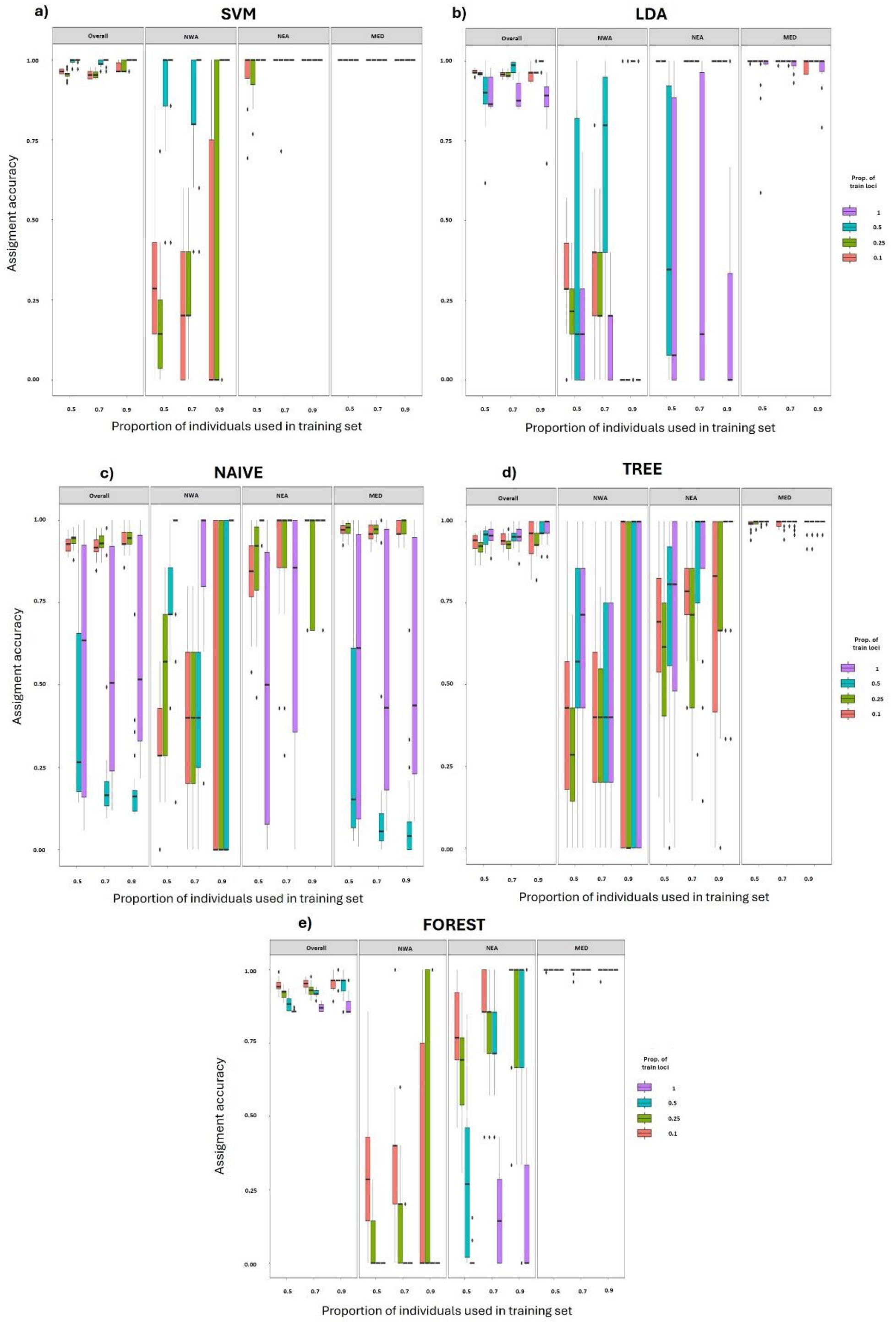


**Figure S5. Results of the five different models implemented in 'assignPOP' package when performing self-assignments at RMU level.** For each one of the models (a-e) we provide the boxplot of the assignment accuracy (in percentage of self-assignments) to the overall dataset and each one of the RMUs. The reassignment accuracy was performed with four different proportions of loci for training (0.1, 0.25, 0.5 and 1) and three different proportions of individuals (0.5, 0.7 and 0.9). We considered all individuals in the baseline (Overall), and the individuals from each RMU independently: North-West Atlantic (NWA), North-East Atlantic (NEA), and the Mediterranean (MED).

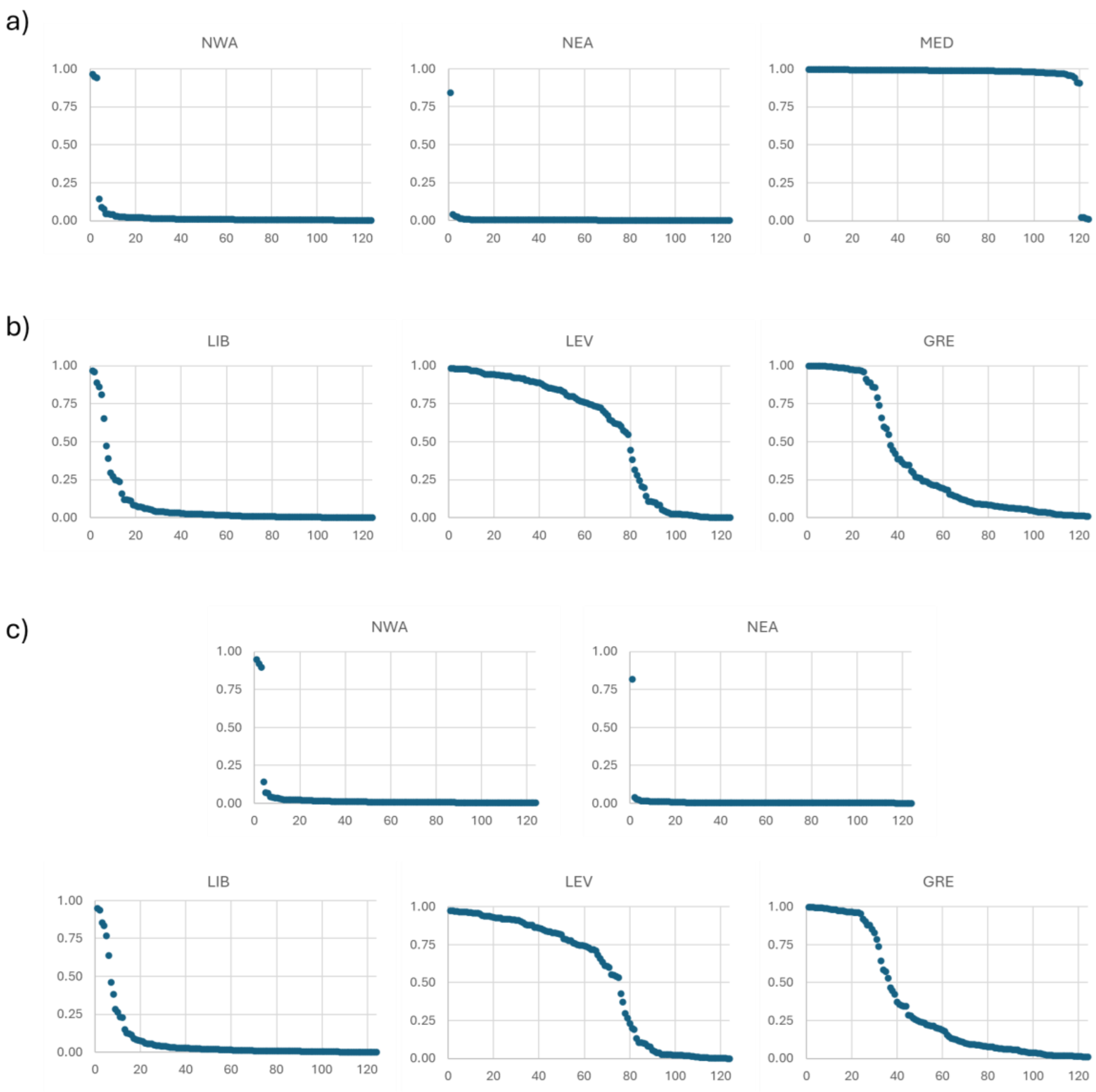


**Figure S6. Individual assignment probabilities of the unknown individual's dataset using different baseline strategies.** a) RMU strategy using the *Caretta Baseline* grouped at the RMU level (NWA, NEA and MED), b) SubRMU strategy using the *MED Baseline* grouped at the SubRMU level (LIB, LEV and GRE), and c) all-at-once strategy using the *Caretta Baseline* grouped in five clusters (NWA, NEA, LIB, LEV and GRE). For each individual the value represented is the mean assignment probability obtained with 10 replicates for each strategy.

**Supplementary table legends.**

Supplementary tables are provided in the Excel file "Supplem_Tables.xlsx"

**Table S1. Nesting sites and foraging ground locations of the 398 *Caretta caretta* individuals genotyped with 2b-RAD.** For each location we provide the group (NES= Nesting or FGR= Foraging),RMU when known (North-West Atlantic (NWA), North-East Atlantic (NEA), and Mediterranean (MED)), including the corresponding SubRMU for the Mediterranean locations (Libya (LIB), Levantine (LEV) and Greece (GRE)), and unknown for FGR (UNK), acronyms (ACR), location latitude and longitude (with a range for FGR), number of individuals (n), mean number of reads per individual (Reads), and mean percentage of missing data (% Missing). * Raw data obtained from Barbanti et al. (2025).

**Table S2. Genomic data of the 398 individuals of *Caretta caretta* based on 6,586 SNPs retained after filtering the Caretta Baseline.** Includes metadata and sequencing statistics for each individual: Group, regional management unit (RMU), location and acronym of the location of the sampling site, unique identifier to each individual (Sample ID), total number of sequencing reads obtained after 2bRAD processing, percentage of reads mapped to the reference genome (% Mapped), proportion of missing data per individual (% Missing), type of indexing used for sequencing, length of the single-end reads generated.

**Table S3. Pairwise FST values (below diagonal) and their significance (above diagonal) based on 6,586 SNPs retained after filtering.** a) Pairwise values between Regional Management Units (NWA, NEA, MED), b) Pairwise values between Mediterranean SubRMUs (LIB, LEV, GRE), and c) Pairwise values between all locations considered in this study (14 locations). Non-significant comparisons after FDR correction are shaded in grey.

**Table S4. Individual assignment of foraging ground samples to Regional Management Units (RMUs) using the 6,586 SNPs from the Caretta Baseline.** For each sample we provide its name (Sample ID), foraging ground sampling location and acronym. For each sample we report the mean probability of assignment to each RMU based on 10 assignment replicates: NWA (North-West Atlantic), NEA (North-East Atlantic), or MED (Mediterranean) and indicate the RMU to which each individual was assigned, based on a probability threshold of >0.75.

**Table S5. Individual assignment of foraging ground samples to SubRegional Management Units (SubRMUs) using the 6,586 SNPs from the MED Baseline.** For each sample we provide its name (Sample ID), foraging ground sampling location and acronym. For each individual we report the mean probability of assignment to each SubRMU based on 10 assignment replicates using the MED Baseline : Libya (LIB), the Levantine region (LEV) and the Greek region (GRE), and indicate the SubRMU to which each individual was assigned, based on a probability threshold of >0.75. Individuals with the maximum assignment probability <0.75 are considered admixed (ADMIX). Note that each individual is listed in the order shown in Table S4.

**Table S6. Individual assignment of foraging ground samples with the all-at-once strategy.** Under this strategy, the individuals of the Caretta Baseline dataset are grouped in five genetic clusters: NWA (North-West Atlantic), NEA (North-East Atlantic), Libya (LIB), the Levantine region (LEV) and the Greek region (GRE). For each sample we provide its name (Sample ID), foraging ground sampling location and acronym. We report the individual mean probability assignment to each of the five subgroups based on 10 replicates and indicate the RMU or SubRMU to which each individual was assigned, based on a probability threshold of >0.75. Individuals with the maximum assignment probability <0.75 are considered admixed (ADMIX). Note that each individual is listed in the order shown in Table S4.
